# Experimental evidence for enhanced receptor binding by rapidly spreading SARS-CoV-2 variants

**DOI:** 10.1101/2021.02.22.432357

**Authors:** Charlie Laffeber, Kelly de Koning, Roland Kanaar, Joyce HG Lebbink

**Affiliations:** Department of Molecular Genetics, Oncode Institute, Erasmus MC Cancer Institute, Erasmus University Medical Center, 3000 CA Rotterdam, The Netherlands; Department of Radiation Oncology, Erasmus University Medical Center, 3000 CA Rotterdam, The Netherlands

## Abstract

Rapidly spreading new variants of SARS-CoV-2 carry multiple mutations in the viral spike protein which attaches to the angiotensin converting enzyme 2 (ACE2) receptor on host cells. Among these mutations are amino acid changes N501Y (lineage B.1.1.7, first identified in the UK), and the combination N501Y, E484K, K417N (B.1.351, first identified in South Africa), all located at the interface on the receptor binding domain (RBD). We experimentally establish that RBD containing the N501Y mutation results in 9-fold stronger binding to the hACE2 receptor than wild type RBD. The E484K mutation does not significantly influence the affinity for the receptor, while K417N attenuates affinity. As a result, RBD from B.1.351 containing all three mutations binds 3-fold stronger to hACE2 than wild type RBD but 3-fold weaker than N501Y. The recently emerging double mutant E484K/N501Y binds as tight as N501Y. The independent evolution of lineages containing mutations with different effects on receptor binding affinity, viral transmission and immune evasion underscores the importance of global viral genome surveillance and functional characterization.

## Main text

Since its emergence in late 2019, SARS-CoV-2 has rapidly spread across the globe, resulting in more than 110 million confirmed COVID-19 cases and more than 2.4 million confirmed casualties as of February 20th, 2021 (coronavirus.jhu.edu). As revealed by the GISAID initiative (gisaid.org), SARS-CoV-2 slowly but continuously mutates, resulting in some instances in variants that become dominant in the population due to increased transmission and/or immune evasion. A number of these variants carry mutations in the spike (S) protein, which is located on the viral surface and interacts with the angiotensin-converting enzyme (ACE2) on host cells, resulting in membrane fusion and viral entry. One such mutation is D614G, which confers increased infectivity and transmissibility and has rapidly become the dominant global variant [1, 2]. Another common mutation is N439K, which is located in the receptor binding domain and enhances affinity for the hACE2 receptor by creating a new salt-bridge across the binding interface [3]. SARS-CoV-2 N439K retains fitness and causes infections with similar clinical outcome, but also shows immune evasion.

In December 2020, new variants of concern have been identified in the UK (B.1.1.7; [4]), South Africa (B.1.351; [5]) and Brazil (P.1 and P.2, both descendants from B.1.1.28; [6, 7]). These strains carry multiple mutations in the spike protein and form the dominant variants in multiple countries. B.1.1.7 contains a change from asparagine to tyrosine at position 501 (N501Y) in the receptor binding motif of the receptor binding domain (RBD). A high-throughput deep mutational scan using yeast predicts this change to increase binding affinity for the hACE2 receptor [8]. Furthermore, N501Y was identified as adaptive mutation during serial passaging of a clinical SARS-CoV-2 isolate in mice [9]. The N501Y mutation is accompanied by two additional changes at the receptor binding interface in the other strains; glutamate to lysine at position 484 (E484K) and lysine to asparagine or threonine at position 417 (K417N in B.1.351; K417T in P.1). P.2 contains only the E484K change in its receptor binding domain. While for B.1.1.7 increased transmission has been established [10], increased prevalence of the other lineages may (also) be due to immune escape [5, 11].

Here we report experimental evidence for the changes in receptor binding affinities due to individual and different combinations of these RBD mutations. Residues N501, E484 and K417 are located relatively far apart on the receptor binding motif (Fig. 1A) and mutations will directly impact molecular interactions across the interface (Fig. 1B). Asparagine 501 in the spike RBD forms a single hydrogen bond across the interface with hACE2 tyrosine 41. This hydrogen bond can no longer be formed in the variants. The new RBD tyrosine 501 now forms a hydrogen bond with hACE2 lysine 353 (Fig. 1C). In addition, the aromatic rings of RBD Y501 and hACE2 Y41 are able to stack onto each other and form favorable van der Waals interactions using their pi-electron orbitals (Fig. 1C). E484K is a charge reversal mutation, resulting in the loss of an ion-pair across the interface with hACE2 lysine 31. A new ion-pair can be formed with neighboring glutamate 35 in the same hACE2 alpha-helix (Fig. 1D), probably accompanied by local rearrangements of flexible side chains to avoid energetically unfavorable electrostatics with lysine 31. According to deep mutational scanning analysis, the polar rearrangements upon mutation of residue 484 have a minor stabilizing effect [8]. The K417N mutation is expected to be destabilizing as replacement of the lysine with a shorter asparagine (in B.1.351, or threonine in P.1) will destroy the salt-bridge across the interface (Fig. 1B). A similar mutation (K417V) was experimentally shown to reduce affinity 2-fold [3].

**Fig. 1.**
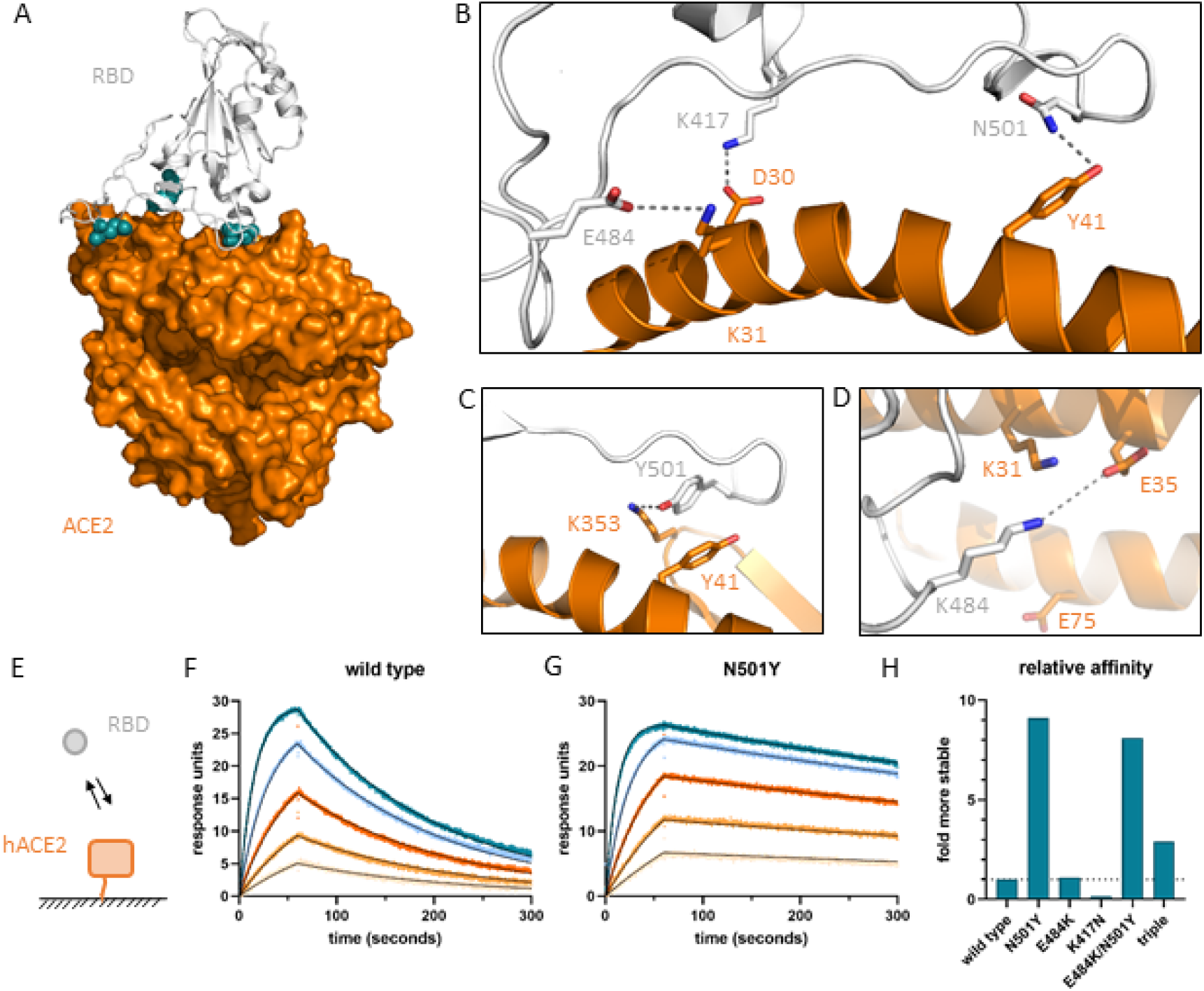
Effects of amino acid mutations on molecular interactions and strength of the interface between hACE2 and the SARS-CoV-2 receptor binding domain. (A) Location of residues N501, E484 and K417 indicated by blue spheres at the interface of SARS-CoV-2 receptor binding domain (white cartoon representation) and the hACE2 ectodomain (orange surface representation) in 6m0j.pdb [24]. (B) Details of the interactions with RBD N501 forming a hydrogen bond with hACE2 Y41, RBD E484 forming an ion-pair with hACE2 K31, and RBD K417 forming a salt-bridge (ion-pair plus hydrogen bond) with hACE2 D30. (C) RBD Y501 creates a new hydrogen bond and stacks its aromatic ring onto hACE2 Y41. (D) Possible new ion-pair between K484 and E35 across the interface. (E) Cartoon visualizing SPR setup using Biacore T100. (F) Sensorgrams for wild type RBD (colored) with fit of a 1:1 binding model (thin black lines). (G) Sensorgrams for N501Y with fit of 1:1 binding model (thin black lines). (H) Affinity of RBD mutants relative to wild type RBD.

To experimentally determine if N501Y indeed increases binding affinity for the hACE2 receptor by itself and in combination with the changes at residues 484 and 417, we purified the hACE2 ectodomain and different SARS-CoV-2 RBD variants from human cells, and determined rate and affinity constants for complex formation using surface plasmon resonance (Fig. 1E-H; supplemental data, fig. S1). Obtained rate and affinity constants for wild type RBD (Fig. 1F; table S1) are similar to previously published values [3]. Strikingly, the increase in affinity for the N501Y variant is 9.1-fold (K_D_ = 1.5 nM instead of 14 nM; Fig. 1G and H, table S1). This is relatively large for a single amino acid change and typical for mutations that improve the hydrophobic effect, in this case probably due to the ring stacking of the two tyrosine side chains across the interface. The change in affinity is predominantly caused by a reduction in the dissociation rate constant, indicating the N501Y spike protein remains bound to the receptor for a longer time period than wild type RBD, increasing the chance to undergo the proper conformational change and induce membrane fusion and cell entry.

The effect of the single E484K mutation on binding affinity is negligible (1.1-fold; Fig. 1H, table S1, fig. S1). Obviously, plasticity at the receptor binding interface allows rearrangements of the polar interactions into a new local conformation that is energetically as favorable as for the original RBD; not only the equilibrium binding constant but also association and dissociation rate constants are similar. The single K417N mutation destabilizes the interaction with hACE2 through a combination of slower binding and faster dissociation (Fig. 1H, table S1, fig. S1).

The combination of all three mutations, as present in strain B.1.351 (first identified in South Africa), results in a 3-fold less stable complex than for N501Y alone due to the effect of K417N, but still 3-fold more stable than with wild type RBD (Fig. 1H, table S1, fig. S1). As the K417T mutation in strain P.1 (first identified in Brazil) will likewise destroy the intramolecular salt bridge, we expect an intermediate affinity for this variant. Double mutant E484K/N501Y has a similar affinity compared with the N501Y single mutant, consistent with the negligible effect of the single E484K mutation (Fig. 1H, table S1, fig. S1).

Taken together, our results show that receptor binding domains from rapidly spreading variants of SARS-CoV-2 bind with increased affinity to the hACE2 receptor and that this is predominantly caused by the N501Y mutation. This is in agreement with recent experimental observations of a similar set of RBD mutations [12, 13]. The importance of receptor binding strength for viral transmission is underscored by the overall strong positive correlation between the stabilizing effect of mutations on receptor binding and their incidence in the population [8, 14]. However, unlike several *in silico* predictions [15-17], we find experimentally that the E484K mutation confers almost no effect on receptor affinity. Mutation of glutamate to lysine at this position abrogates binding to certain antibodies and results in immune evasion, reinfection and reduced efficacy of vaccines [11, 18-22]. It is therefore possible that the increased prevalence of lineage P.2, carrying only E484K in its RBD, is due to immune escape rather than increased transmissibility. The recently reported observation of the independent emergence of the E484K mutation into the more transmissible B.1.1.7 strain in for example the UK [19] is of particular concern, as it combines the immune evasion properties of the E484K mutation with N501Y’s high affinity, as shown here for the E484K/N501Y double mutant. Additional adaptation seems to have occurred in certain global regions with relatively high previous exposure; mutation of K417 in P.1 and B.1.351 confers additional immune evasion [18] at the expense of RBD-receptor complex stability due to loss of the salt bridge across the interface. The K417N mutation in B.1.351 has occurred as an independent event onto the E484K/N501Y combination before rapidly spreading into the population [5]. As herd immunity will build up due to increased exposure and vaccination, continuous genomic and functional characterization will be crucial to tailor restrictive guidelines, vaccine composition and vaccination strategies towards optimal control of variants with increased receptor affinity, transmissibility and/or immune evasion.

## Materials and Methods

### Construction of expression clones

The hACE2 ectodomain (residues 18-615) was fused with the HA signal sequence, a linker containing a 8*HIS-tag, a Twin-Strep-tag and TEV protease cleavage site into vector pCEP4 using Gibson assembly [23]. The same strategy was used to create the expression plasmid for wild type SARS-CoV-2 spike receptor binding domain (reference genome Wuhan-HU-1; residues 333-529). Templates used to create the PCR fragments were pTwist-EF1alpha-SARS-CoV-2-S-2xStrep (gift from Nevan Krogan; Addgene plasmid #141382), pCEP4-myc-ACE2 (gift from Erik Procko; Addgene plasmid #141185). Oligonucleotides for Gibson assembly and site-directed mutagenesis were obtained from IDT (Integrated DNA Technologies, Coralville, USA). Single amino acid changes in the RBD (N501Y, E484K, K417N) and combinations thereof were introduced using QuikChange (XL Site-Directed Mutagenesis Kit, Agilent, Santa Clara, USA). Resulting expression plasmids were purified using Nucleobond Xtra maxi, (Macherey-Nagel, Dueren, Germany) and sequence verified.

### Cell culture

Human embryonic kidney 393T cells were adapted to suspension culture in Gibco Freestyle 293 medium (Thermo Fisher Scientific, Waltham USA) supplemented with 20 U/L penicillin, 100 mg/L streptomycin (Sigma Aldrich, Saint Louis USA) and 1% FCS (Capricorn scientific, Ebsdorfergrund Germany) at 37°C, 5% CO_2_, 150 RPM. Cell densities were kept between 0.3 -3 *10^6^ cells/ml during propagation. For transfections, cells were seeded to a density of 0.7 *10^6^ cells/ml and incubated for 24 hours. Plasmid DNA (1 mg) was mixed by dropwise addition of 2 mg linear polyethylenimine M_W_ 25000 (Polysciences Inc, Warrington USA) in 25 ml Gibco Hybridoma-SFM (Thermo Fisher Scientific, Waltham USA), incubated for 30 minutes, added to HEK293T cells (25 ml/1 liter cell culture) and incubated for 72 hours.

### Protein purification

Cell culture medium was cleared by centrifugation at 1000 g for 15 minutes and imidazole was added to a final concentration of 20 mM. Medium was stirred for 1 hour at 4°C in the presence of 5 ml Ni-NTA Agarose beads (Qiagen, Venlo, The Netherlands) washed with 25 mM Hepes-KOH pH 7.5, 150 mM NaCl (buffer A). The beads were collected in a gravity flow column, and washed 3 times with buffer A supplemented with 20 mM imidazole. Protein was eluted with buffer A supplemented with 250 mM imidazole. For hACE2, eluate was loaded on a 1 ml StrepTrap HP column (Cytiva Marlborough USA) operated by an ÄKTA pure (Cytiva, Marlborough USA). hACE2 was eluted with 2.5 mM desthiobiotin in buffer A. Eluate containing RBD variants was incubated with 1 ml Streptactin XT superflow beads (IBA GmbH, Göttingen, Germany) for 1 hour at 4°C. Beads were collected by centrifugation for 1 minute at 200 g, washed 3 times with buffer A and incubated over night at 4°C in the presence of 0.3 mg reducing-agent free TEV protease. Eluate was supplemented with 20 mM imidazole and TEV protease was removed using Ni-NTA agarose beads. hACE2 and RBD eluates were buffer exchanged against buffer A using a 5ml HiTrap desalting column (Cytiva, Marlborough, USA) operated by ÄKTA pure. Aliquots were snap frozen in liquid nitrogen and stored at -80°C. Protein concentrations were determined using OD^280 nm^ measured on a Nanodrop 2000 (Thermo Scientific, Waltham USA) and extinction coefficients of ε _m_ ^280 nm^ = 101,170 M^-1^cm^-1^ for tagged hACE2, ε _m_ ^280 nm^ = 33,850 M^-1^cm^-1^ for wild type, E484K and K417N RBD, and ε _m_^280 nm^ = 35,340 M^-1^cm^-1^ for N501Y, double and triple mutant RBD.

### Surface Plasmon Resonance

Surface plasmon resonance (SPR) spectrometry was performed at 25°C on a Biacore T100 (Cytiva, Marlborough USA). A Series S CM5 sensor chip surface was derivatized with 1500 response units (RU) of Streptactin XT (Twin-Strep-tag® Capture Kit; IBA GmbH, Göttingen, Germany) using the amine coupling kit (Cytiva, Marlborough USA). Flow cell 1 was used for reference subtraction, flow cell 2 was derivatized with 100 RU hACE2 using the capture procedure. Increasing concentrations of RBD variants (0-125 nM) in 25 mM HEPES-KOH pH 7.5, 150 mM NaCl and 0.05% Tween 20 were injected across the chip at 50 µl/min. Flow cells were regenerated using 3 M GuHCl (Sigma Aldrich, Saint Louis USA). Sensorgrams were analysed, fit with a 1:1 binding model using BiaEvaluation software (Cytiva, Marlborough USA), and visualized using Prism (GraphPad Software, San Diego, USA).

### Structure analysis

An optimized model for the complex between hACE2 receptor and the SARS-CoV-2 receptor binding domain was created from 6m0j.pdb [24] using PDB-redo [25]. This resulted, amongst others, in a side chain flip for asparagine 501 in the RBD such that it forms a hydrogen bond across the complex interface with Tyrosine 41 in ACE2. Models for mutant RBDs N501 and E484K were created using Misssense3D [26]. Models were analyzed and figures were created using PyMOL (The PyMOL Molecular Graphics System, Version 2.0 Schrödinger, LLC).

## Acknowledgments

We are grateful to Marion Koopmans for valuable comments and critical reading of the manuscript. Funding: this work is supported by the gravitation program CancerGenomiCs.nl from the Netherlands Organisation for Scientific Research (NWO), part of the Oncode Institute, which is partly financed by the Dutch Cancer Society, and NWO grant OCENW.XS.055. Author contributions: R.K and J.L supervised the project. K.K, C.L and J.L created expression clones. K.K and C.L purified proteins. C.L. performed SPR analysis. C.L and J.L. analyzed SPR data. J.L performed structure analysis and visualized results. C.L and J.L wrote draft, C.L, J.L and R.K edited manuscript. Competing interests: Authors declare no competing interests.

---

**Figure S1:**
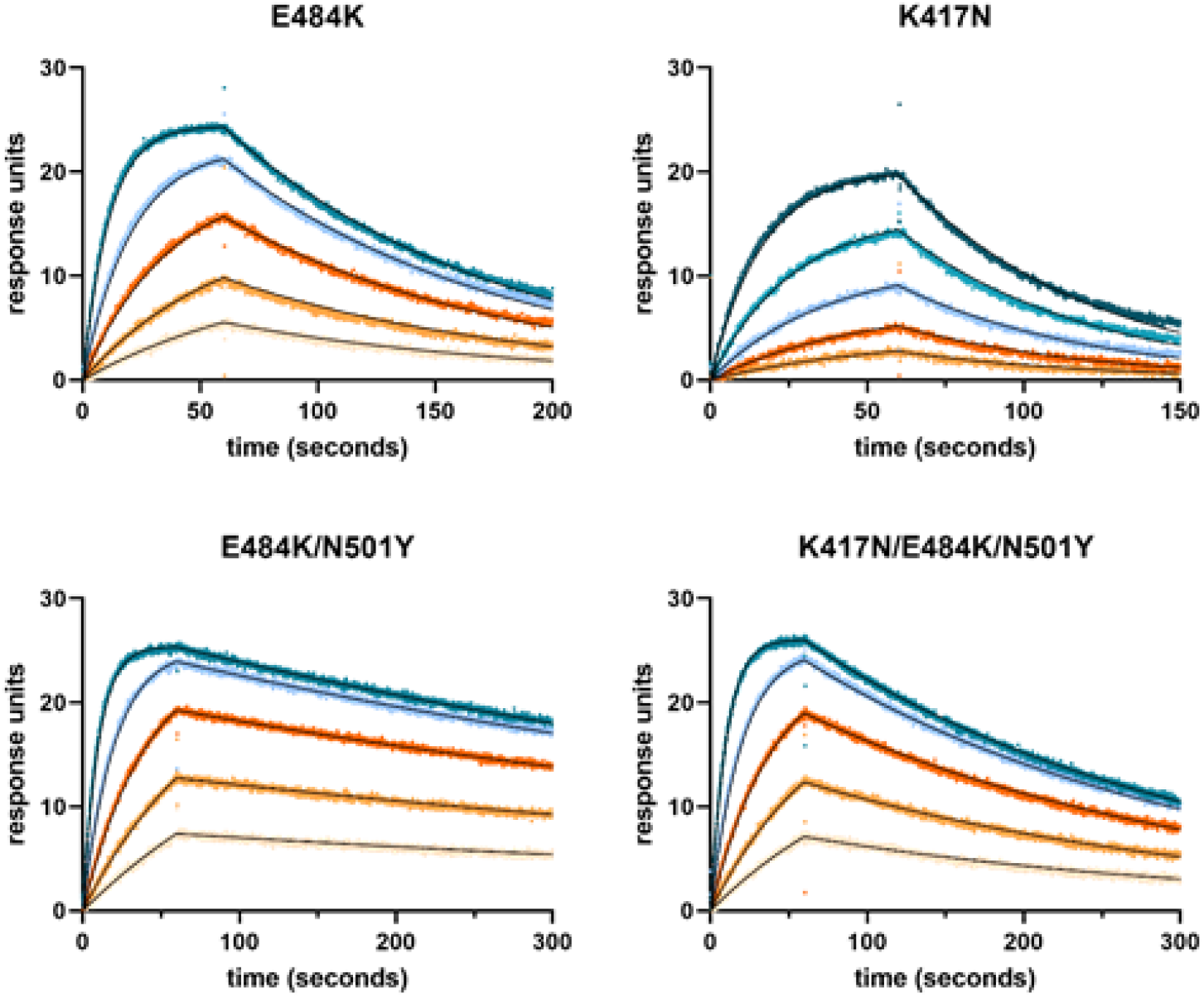
SPR sensorgrams with fit of 1:1 binding model for different RBD variants.

**Table S1.**
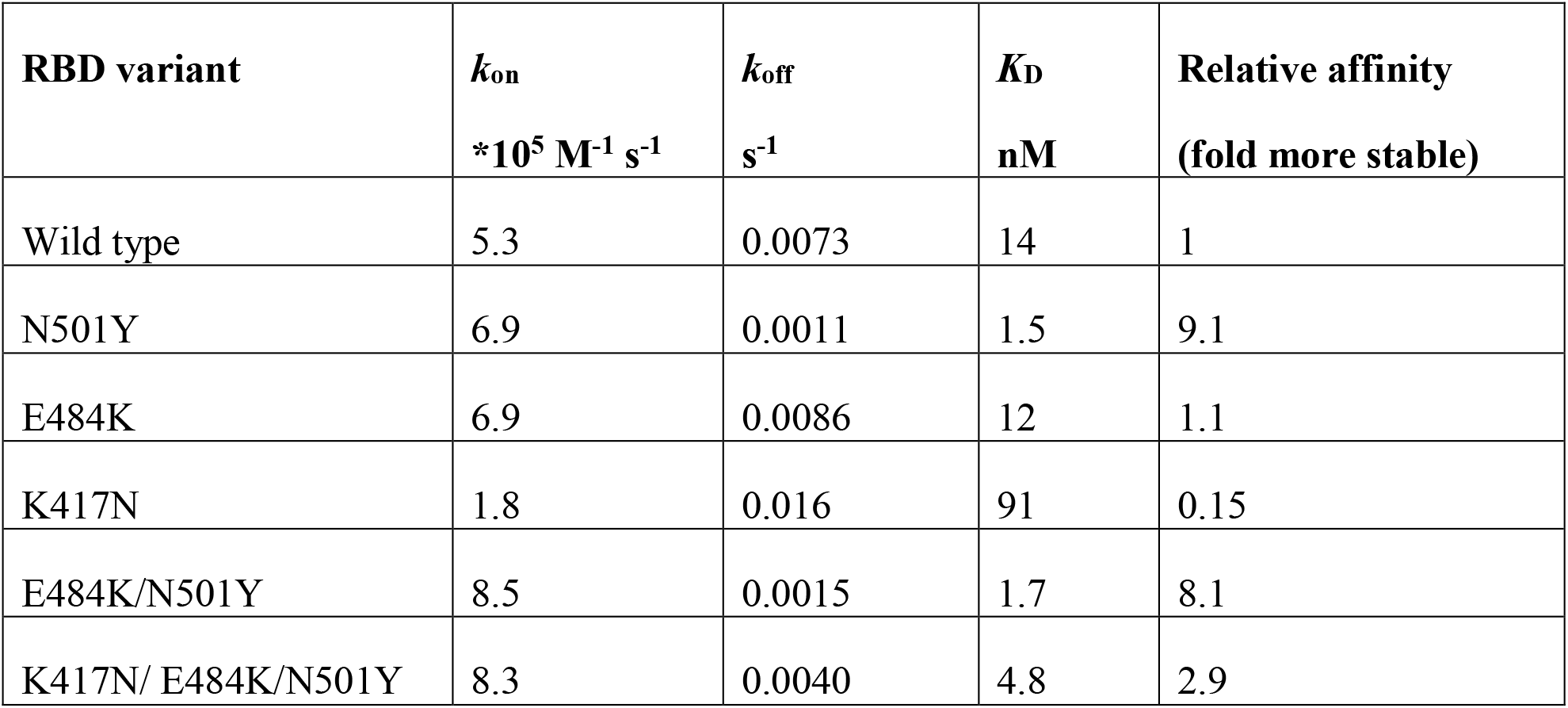
Rate and affinity constants for binding of variants of the SARS-CoV-2 RBD to the human ACE2 ectodomain determined using surface plasmon resonance. Relative affinity of the mutant variants corresponds to the fold decrease in equilibrium binding constant relative to wild type.

